# Sequence characterization of eccDNA content in glyphosate sensitive and resistant Palmer amaranth from geographically distant populations

**DOI:** 10.1101/2021.11.19.469340

**Authors:** Hailey Spier Camposano, William T. Molin, Christopher A. Saski

## Abstract

The discovery of non-chromosomal circular DNA offers new directions in linking genome structure with function in plant biology. Glyphosate resistance through *EPSPS* gene copy amplification in Palmer amaranth was due to an autonomously replicating extra-chromosomal circular DNA mechanism (eccDNA). CIDER-Seq analysis of geographically distant glyphosate sensitive (GS) and resistant (GR) Palmer Amaranth (*Amaranthus palmeri*) revealed the presence of numerous small extra-chromosomal circular DNAs varying in size and with degrees of repetitive content, coding sequence, and motifs associated with autonomous replication. In GS biotypes, only a small portion of these aligned to the 399 kb eccDNA replicon, the vehicle underlying gene amplification and genetic resistance to the herbicide glyphosate. The aligned eccDNAs from GS were separated from one another by large gaps in sequence. In GR biotypes, the eccDNAs were present in both abundance and diversity to assemble into a nearly complete eccDNA replicon. Mean sizes of eccDNAs were similar in both biotypes and were around 5kb with larger eccDNAs near 25kb. Gene content for eccDNAs ranged from 0 to 3 with functions that include ribosomal proteins, transport, metabolism, and general stress response genetic elements. Repeat content among smaller eccDNAs indicate a potential for recombination into larger structures. Genomic hotspots were also identified in the Palmer amaranth genome with a disposition for gene focal amplifications as eccDNA. The presence of eccDNA may serve as a reservoir of genetic heterogeneity in this species and may be functionally important for survival.

## Introduction

Extra-chromosomal circular DNA (eccDNA) are nucleus limited ring-like DNA entities derived from the genome and have been found in a wide range of eukaryotic organisms including yeast, *Drosophila*, *Xenopus*, mice, and humans [1–4]. In yeast, eccDNAs with functional genes and sizes of up to 38 kb that cover 23% of the genome have been reported [5]. EccDNAs have been reported in normal healthy cells in humans [6, 7] with functions associated with aging and the formation of telomeric circles [8, 9], cancer progression, and therapeutic resistance [10–12]. EccDNAs have been implicated in approximately half of all human cancers contributing to genetic heterogeneity that enables aggressive tumors with a selective advantage; hence the higher prevalence in malignant tumors [13–15]. Sizes of cancer related eccDNA have been reported to range from several hundred base pairs to 600 kb encoded with functional oncogenes and their various regulatory elements [16, 17]. In plants, eccDNAs have been reported in *Arabidopsis* [18, 19], *Oryza*, *Pisum*, *Secale*, *Triticum*, and *Vicia* [20, 21] with sizes that range from 1 kb to 50kb. These eccDNAs contain coding sequences commonly found within the nucleus such as ribosomal genes, tRNAs, and transposons [19, 22, 23]. EccDNAs are thought to arise from linear chromosomes through repeat-mediated intrachromosomal homologous recombination that results in the ‘looping-out’ of circular structures. These focal amplifications are mediated by multimers corresponding to 5S ribosomal DNA, non-coding chromosomal high-copy tandem repeats, and telomeric DNA [1, 21, 22]. In *Arabidopsis*, eccDNA genesis is the result of recombination among inverted repeats upstream and downstream of the various tRNAs and transposons [19]. Several follow up studies using *Arabidopsis* and rice, have shown that defective RNA polymerase II (Pol II) activity and simultaneous inhibition of DNA methylation leads to the activation of retrotransposons which can induce eccDNA formation upon stress [11]. These studies suggest a possible relationship among epigenetic status, regulation of transposon bursts, and genomic focal amplifications.

The presence of eccDNA with functional genes in a cell can be a signature of a stress and/or function as a reservoir of genetic variation in which a cell may activate as a rapid response to stress. For example, oncogene amplification and expression via eccDNA in human cancers provides a unique mechanism for massive gene expression [24] and ultimately a reservoir of genetic heterogeneity by which cancer cells have a selective advantage for aggressive behavior and persistence [13].

Recently, the genetic entity conferring resistance to the herbicide glyphosate in Palmer amaranth (*Amaranthus palmeri*), now termed the eccDNA replicon, was revealed to be a massive, 399 kb extrachromosomal circular DNA (eccDNA) [25, 26]. Glyphosate resistance in Palmer amaranth is achieved through replicon amplification with simultaneous gene copy amplification and expression of the 5-enoylpyruvylshikimate-3-phosphate synthase (*EPSPS*) gene and its product, EPSP synthase [27], which is the herbicide target of glyphosate [28]. Glyphosate resistance may occur with as few as 5 copies of *EPSPS.* The increase in *EPSPS* functions to ameliorate the unbalanced or unregulated metabolic changes, such as shikimate accumulation, loss of aromatic amino acids, phenolic acids for lignin synthesis, and structural intermediates for plant growth regulators associated with glyphosate activity in sensitive plants [27, 29]. Isolation and single-molecule sequencing of the replicon resulted in a single copy of the *EPSPS* gene along with 58 other predicted genes whose broad functions traverse detoxification, replication, recombination, DNA binding, and transport [26, 30]. Gene expression profiling of the replicon under glyphosate treatment showed transcription of 41 of the 59 genes in GR biotypes, with high expression of *EPSPS*, aminotransferase, zinc-finger, and several uncharacterized proteins [26, 30].

Repeat sequences and mobile genetic elements have been associated with eccDNA formation [4, 6, 18, 20, 30, 31] in higher eukaryotes. The repeat landscape of the replicon is described as a complex arrangement of repeat sequences and mobile genetic elements interspersed among arrays of clustered palindromes which may function in stability, DNA duplication and/or a means of nuclear integration [26]. In a follow up study, sequence analysis identified a region in the replicon with elevated A+T content and an exact match to a conserved eukaryotic extended autonomous consensus sequence (EACS) [32]. Surrounding this sequence were multiple DNA unwinding elements (DUE), which together are often associated with DNA bending and origins of replication and typically found near EACS [33, 34]. Regions flanking these elements in the replicon were cloned into an ARS-less yeast plasmid which resulted in colony formation, suggesting autonomous replication as the mechanism for the replicon increases in copy number [35].

Initial low-resolution FISH analysis of GR *A. palmeri* showed the amplified *EPSPS* gene was randomly distributed in the genome, suggesting a possible transposon-based mechanism of mobility [27]. A follow up study using much longer bacterial artificial chromosome (BAC) probes coupled with high resolution fiber extension microscopy verified the eccDNA replicon and identified various structural polymorphisms including intact, circular, dimerized circular, and linear forms [25]. Additionally, this study resolved a critical question regarding the maintenance mechanism that explains uneven segregation of glyphosate resistance among progenies – genomic tethering. Analysis of fiber-FISH images with replicon probes and meiotic pachytene chromosomes revealed very clear, single signals [25]. If the replicon were integrated into the genome, then double signals would be evident, suggesting a tethering mechanism as a means of genomic persistence to daughter cells during cell division [25]. Other genetic entities that maintain genomic persistence through tethering include DNA viruses such as Epstein-Barr, Rhadinovirus, Papillomavirus, and others [36].

Glyphosate resistance in Palmer amaranth has been observed in individuals with *EPSPS* copy numbers that range from 5-150 copies [27, 37, 38]. Amplification of the *EPSPS* gene correlated with amplification of flanking genes and sequence [26, 30], which suggests a large amplification unit and genome size enlargement in cells with many replicon copies [30]. Flow cytometry verified significant genome expansion in plants with high copy numbers (eg. 11% increase in genome size with ∼100 extra copies of the replicon), seemingly without fitness penalty [30].

Glyphosate resistance in Palmer amaranth was originally reported in Georgia in the early 2,000’s [39], and a recent analysis using whole genome shotgun sequencing verified that the replicon was present and intact in GR Palmer amaranth populations across the USA [40, 41]. This study also reported a lack of replicon SNP variation among GR eccDNAs from geographically distant states when aligned to the Mississippi replicon reference [26]. The replicon was not present in GS individuals, which supports a single origin hypothesis and spread of the replicon across the USA through mechanical means such as spread of GR pollen in contaminated plant products, on farm equipment, and cattle movement, or via pollen.

The genomic mechanisms, origins and how the replicon assembled and gave rise to eccDNA in Palmer amaranth remains elusive, but the above studies lead to a couple of hypotheses: 1) the eccDNA replicon formed through intramolecular recombination among distal parts of the nuclear genome in short evolutionary time, or 2) there may exist a reservoir of smaller eccDNAs that are basal in the cell that may have the ability to recombine to assemble larger units as part of a dynamic response to stress. In this study, we report the presence and sequence characterization of an abundant reservoir of eccDNAs in both GS and GR biotypes using single molecule sequencing and the CIDER-Seq approach [18]. We examine the similarities and differences among samples representing distant geographic locations reported in [41], quantitate their abundance and diversity and assess whether recombination may be possible to form larger multimeric units.

## Results

### EccDNA content and coding structure in geographically distributed *A. palmeri*

Following the general methods and recommended computational pipelines outlined in the CIDER-Seq single-molecule approach [42], we identified an extensive amount of variable-sized eccDNA in all samples of (GS) and GR) biotypes that were sequenced [Table 1]. The number of unique eccDNAs detected in GS samples ranged from 443 (ks_s) to 6,227 (ms_s) with a mean of 2,661 [Table 1]. Unique eccDNAs were in much higher abundance in GR samples and ranged from 2,200 (az_r) to 5,650 (ms_r), with a mean of 4,448, nearly double that of GS [Table 1]. Length distributions of eccDNA were similar among both GS and GR biotypes and ranged from 27bp to nearly 27kb, with mean lengths of around 6kb, [Table 1 and Fig 1]. Gene prediction resulted in eccDNAs both with and without complete open reading frames. In GS samples, the number of eccDNAs with predicted genes ranged from 76-505 with a mean of 272 eccDNAs with genes per sample. GR eccDNAs with predicted genes was nearly 4 times greater with a range of 263-1,179 and a mean of 718 eccDNA with genes per sample, suggesting that glyphosate stress influenced unique gene focal amplifications [Table 1]. Of the eccDNA with predicted genes, the number of predicted genes per eccDNA ranged from 1 to 10, with an average of 2 genes per eccDNA in both GS and GR [S1 and S2 Tables]. Transfer RNAs (tRNA) were predicted exclusively on eccDNA without CDS sequences and ranged widely from 46-715 (average of 350 per sample) in GS and samples and 130-528 (average of 364 per sample).

**Table 1.**
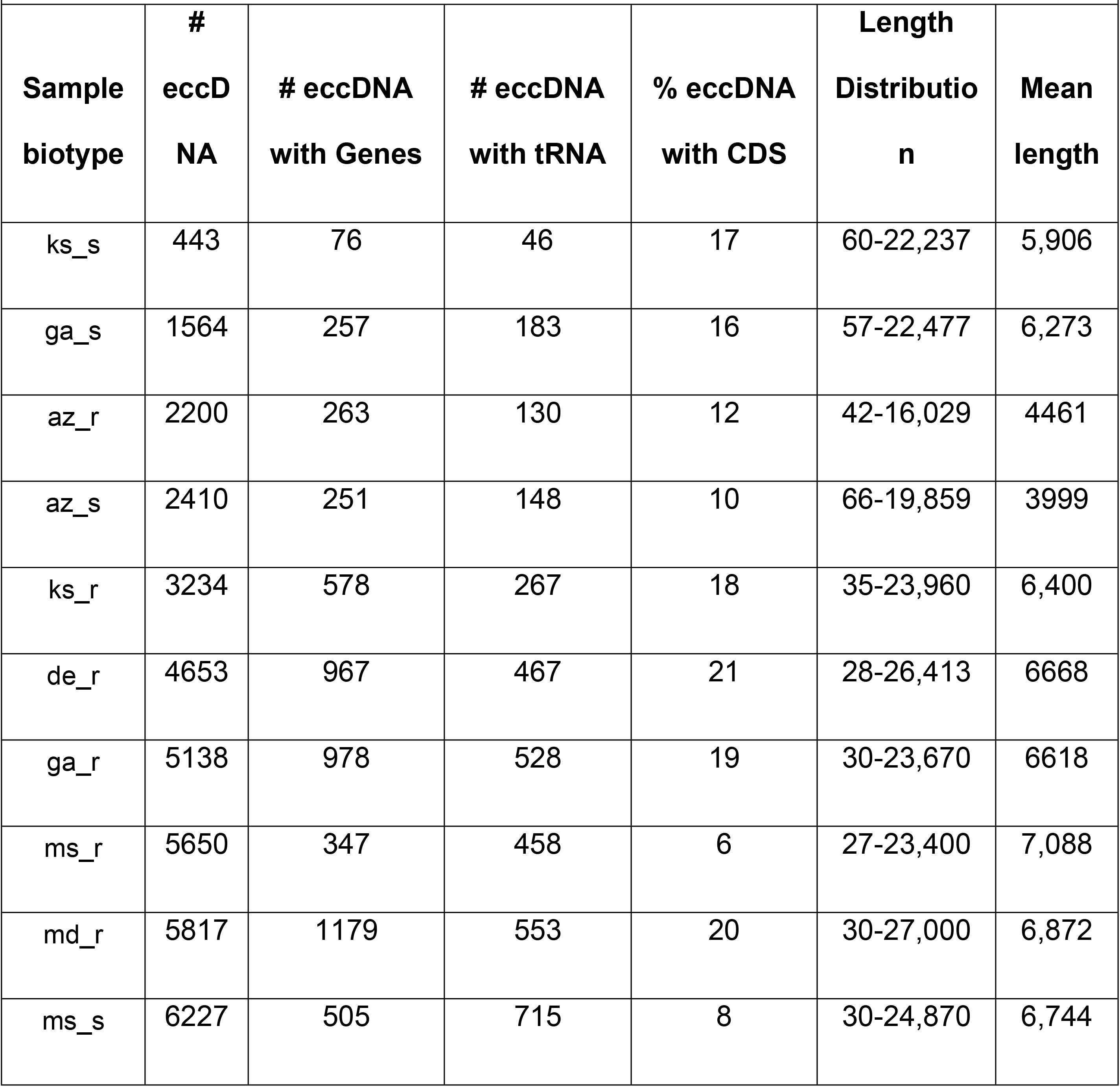
EccDNA characterization of GS and GR biotypes.

**Fig 1.**
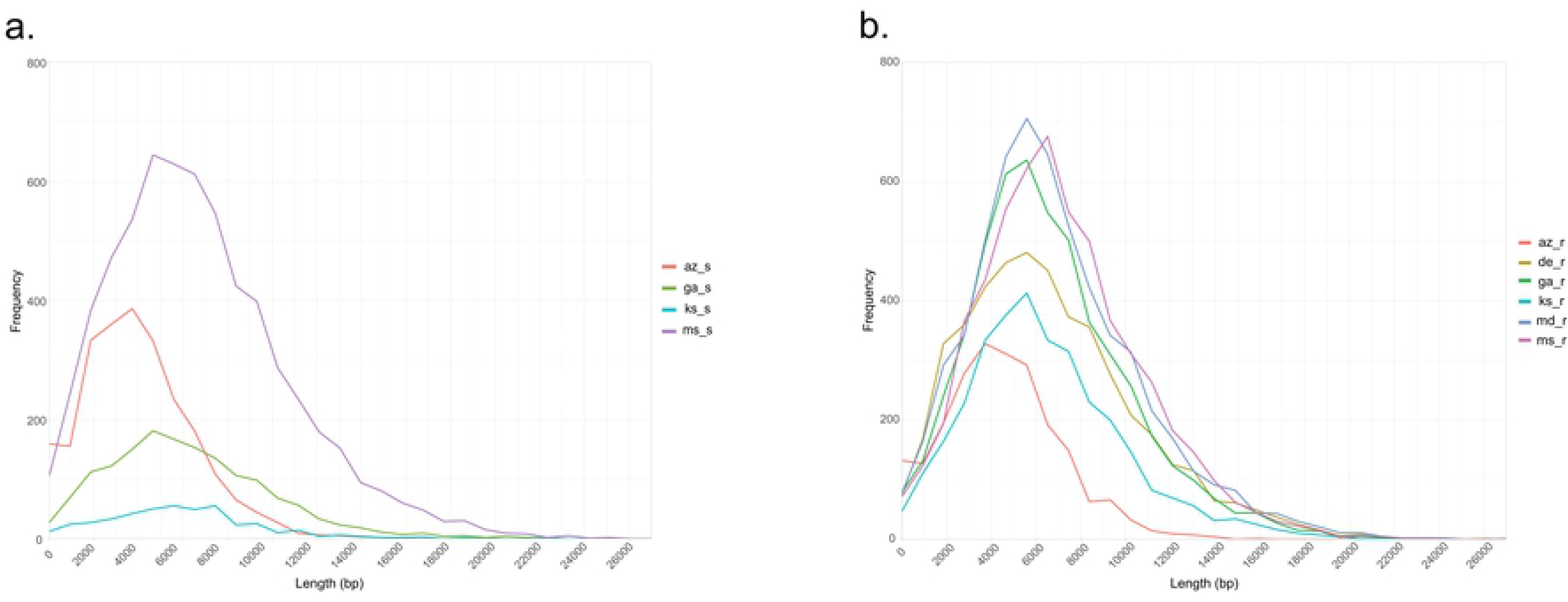
Frequency polygon graph for Lengths (bp) of *A. palmeri* eccDNAs. A) Glyphosate sensitive samples were sourced from Arizona (az_s), Georgia (ga_s), Kansas (ks_s), and Mississippi (ms_s) A. palmeri plants. B) Glyphosate resistant samples were sourced from Arizona (az_r), Delaware (de_r), Georgia (ga_r), Kansas (ks_r), Maryland (md_r), and Mississippi (ms_r) plants.

### Coding content of eccDNAs in glyphosate sensitive and resistant *A. palmeri*

Gene content from both GS and GR biotypes was compared to identify unique and common functional protein coding domains among the geographically distant samples. In GS biotypes, 9 functional protein coding domains were discovered that are common among the each of the states [Fig 2]. These functional domains are annotated as ATP synthase, cytochrome P450, protein kinase, ribosomal protein, NADH dehydrogenase, Clp protease, and oxidoreductase [Table 2]. Various pairwise combinations of GS *A. palmeri* biotypes shared a range of 1 to 12 elements [Fig 2 and S3 Table]. Genes that regulate cell division, such as the Ras protein family and those involved in DNA replication (helicase) were common among Arizona, Georgia, and Mississippi GS eccDNA samples [S3 Table].

**Fig 2.**
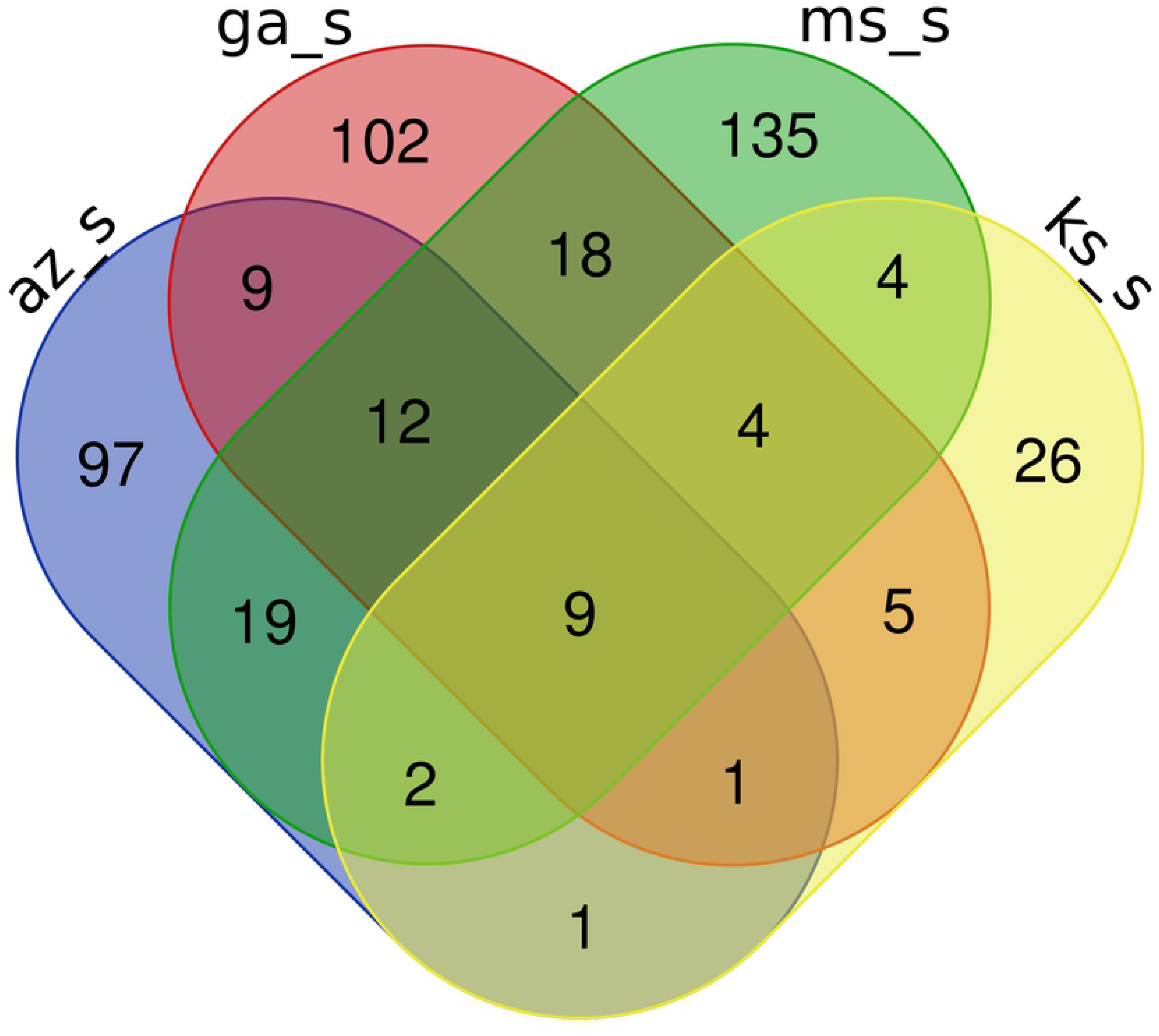
Venn diagram of PFAM elements shared by GS eccDNA samples. Arizona (az_s), Georgia (ga_s), Kansas (ks_s), and Mississippi (ms_s) sensitive A. palmeri eccDNA shared 9 total

**Table 2.**
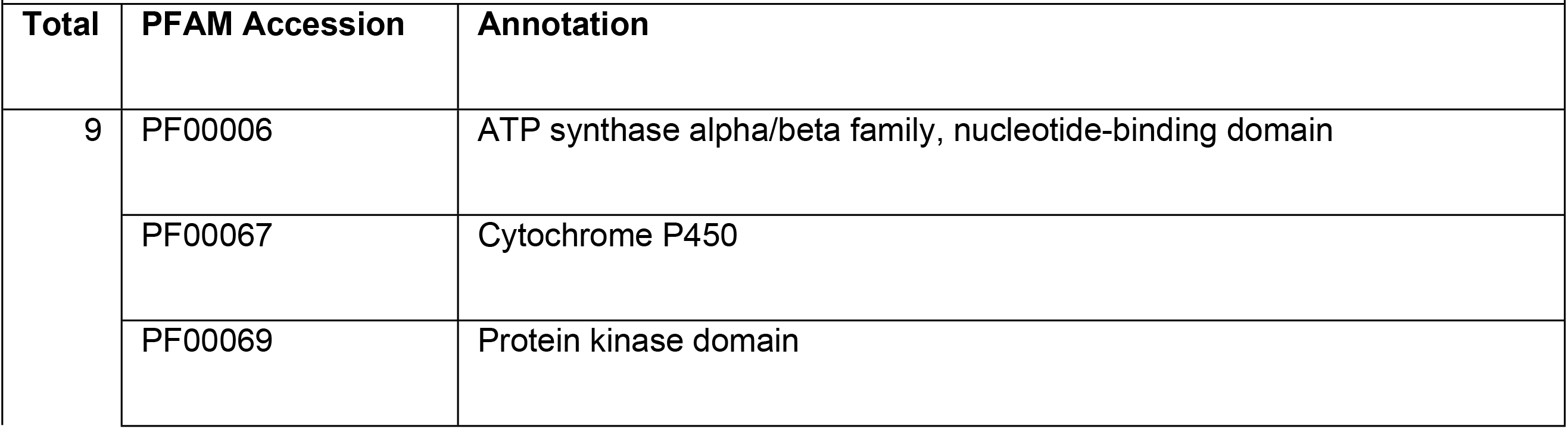
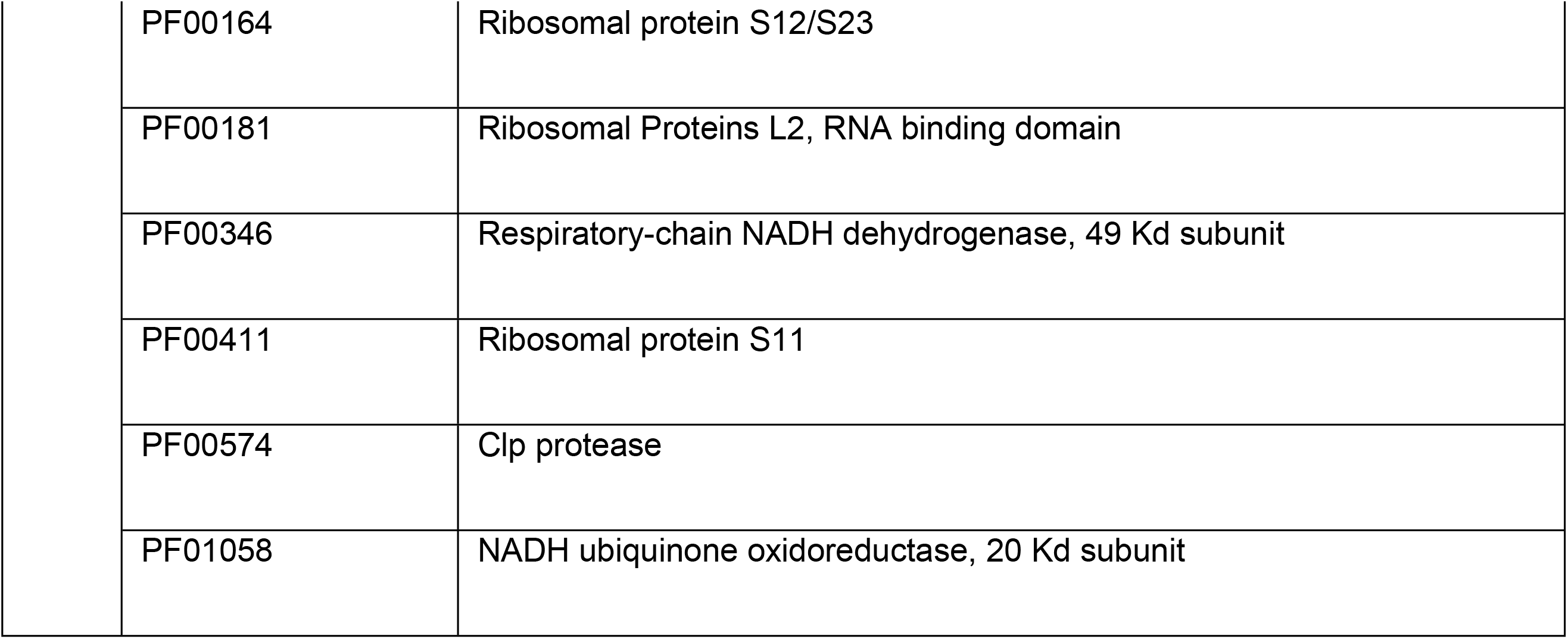
Gene elements shared by all sensitive eccDNA samples.

| Table 2. Gene elements shared by all sensitive eccDNA samples. |  |  |
| --- | --- | --- |
| Total | PFAM Accession | Annotation |
| 9 | PF00006 | ATP synthase alpha/beta family, nucleotide-binding domain |
|  | PF00067 | Cytochrome P450 |
|  | PF00069 | Protein kinase domain |

|  |  |
| --- | --- |
| PF00164 | Ribosomal protein S12/S23 |
| PF00181 | Ribosomal Proteins L2, RNA binding domain |
| PF00346 | Respiratory-chain NADH dehydrogenase, 49 Kd subunit |
| PF00411 | Ribosomal protein S11 |
| PF00574 | Clp protease |
| PF01058 | NADH ubiquinone oxidoreductase, 20 Kd subunit |

Several abiotic/biotic resilience-related functional protein domains were found in Arizona and Mississippi GS samples that includes an oxysterol-binding protein, pectinesterase, NmrA-like family, and WRKY DNA-binding domain elements [S3 Table]. Also discovered were shared functional domains involved in DNA methylation and histone maintenance (H2A/H2B/H3/H4) [S3 Table]. Common between Georgia and Mississippi GS biotypes were ABC transporter and Cytochrome C oxidase subunit II (periplasmic domain) protein domains [S3 Table]. Unique to Arizona were response regulators such as trehalose-phosphatase, chalcone-flavanone isomerase, O-methyltransferase, Myb-like DNA-binding domain [S3 Table]. Hundreds of other unique functional domains in different GS biotypes were recorded in S3 Table. It is notable that the *EPSPS* gene was not found in any of the GS eccDNAs.

In GR biotypes, we identified a total of 20 functional protein domains that are shared among all 6 resistant samples [Fig 3 and Table 3]. The shared GR domains had various cellular maintenance functions in addition to stress response domains that include ABC transporter, HSP70 protein, Ribosomal protein, WD domain, and Leucine rich repeats [Table 3]. A range of 1 to 9 protein family domains were shared by at least 5 of the GR biotypes [Fig 3 and S4 Table]. No apical meristem (NAM) protein, peroxidase, TCP-1/cpn60 chaperonin family are among the stress response elements. Arizona, Delaware, Kansas, and Maryland GR biotypes all contained EPSP synthase (3-phosphoshikimate 1-carboxyvinyltransferase) and Arabidopsis phospholipase-like protein (PEARLI 4) functional domains, with 21 and 24 copies distributed across various eccDNA within these four samples respectively.

**Fig 3.**
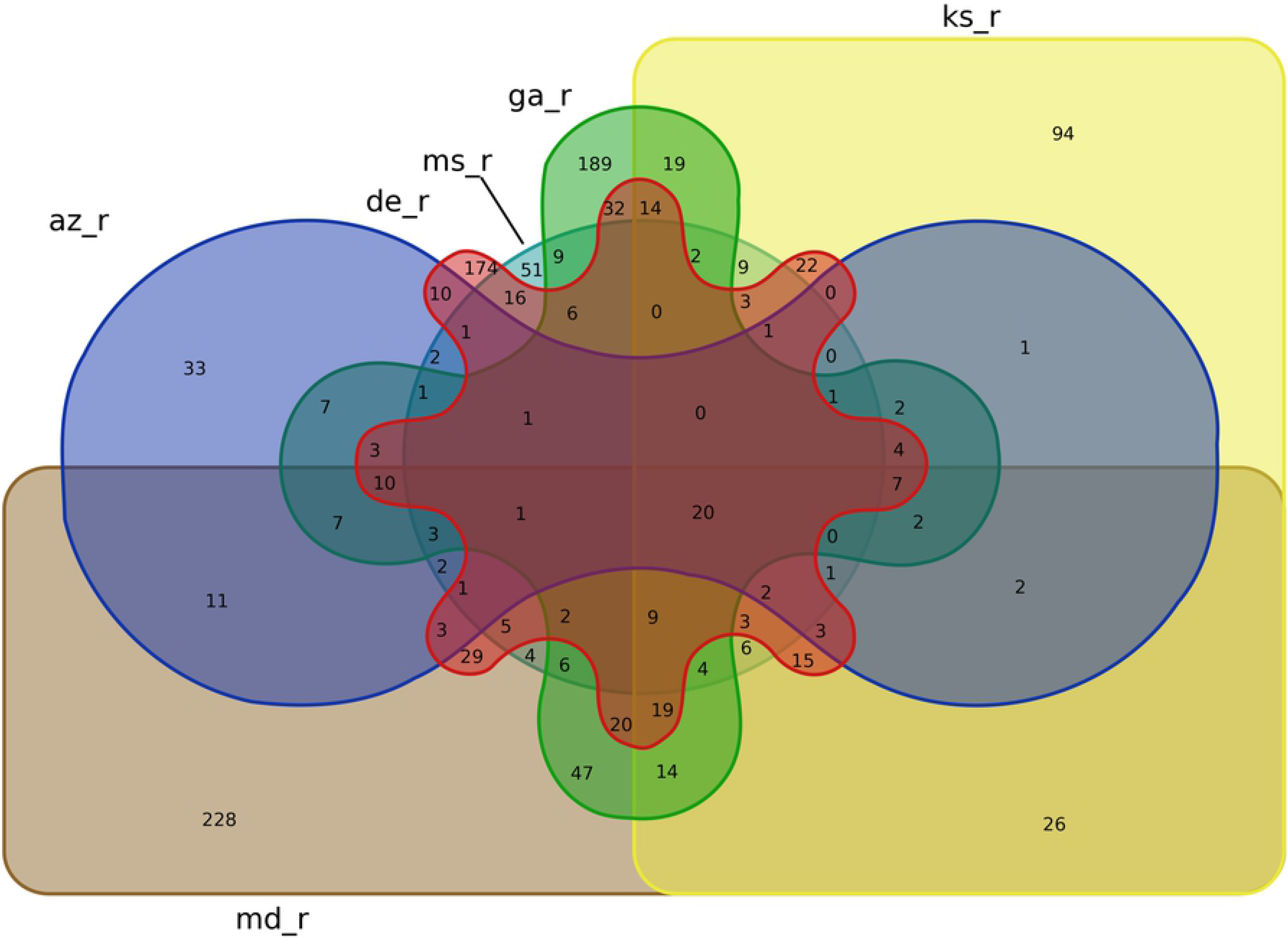
Venn diagram of PFAM elements shared among GR *A. palmeri* eccDNA samples. Arizona (az_r), Delaware (de_r), Georgia (ga_r), Kansas (ks_r), Maryland (md_r), and Mississippi (ms_r) resistant A. palmeri eccDNA shared 20 total PFAM elements.

**Table 3.**
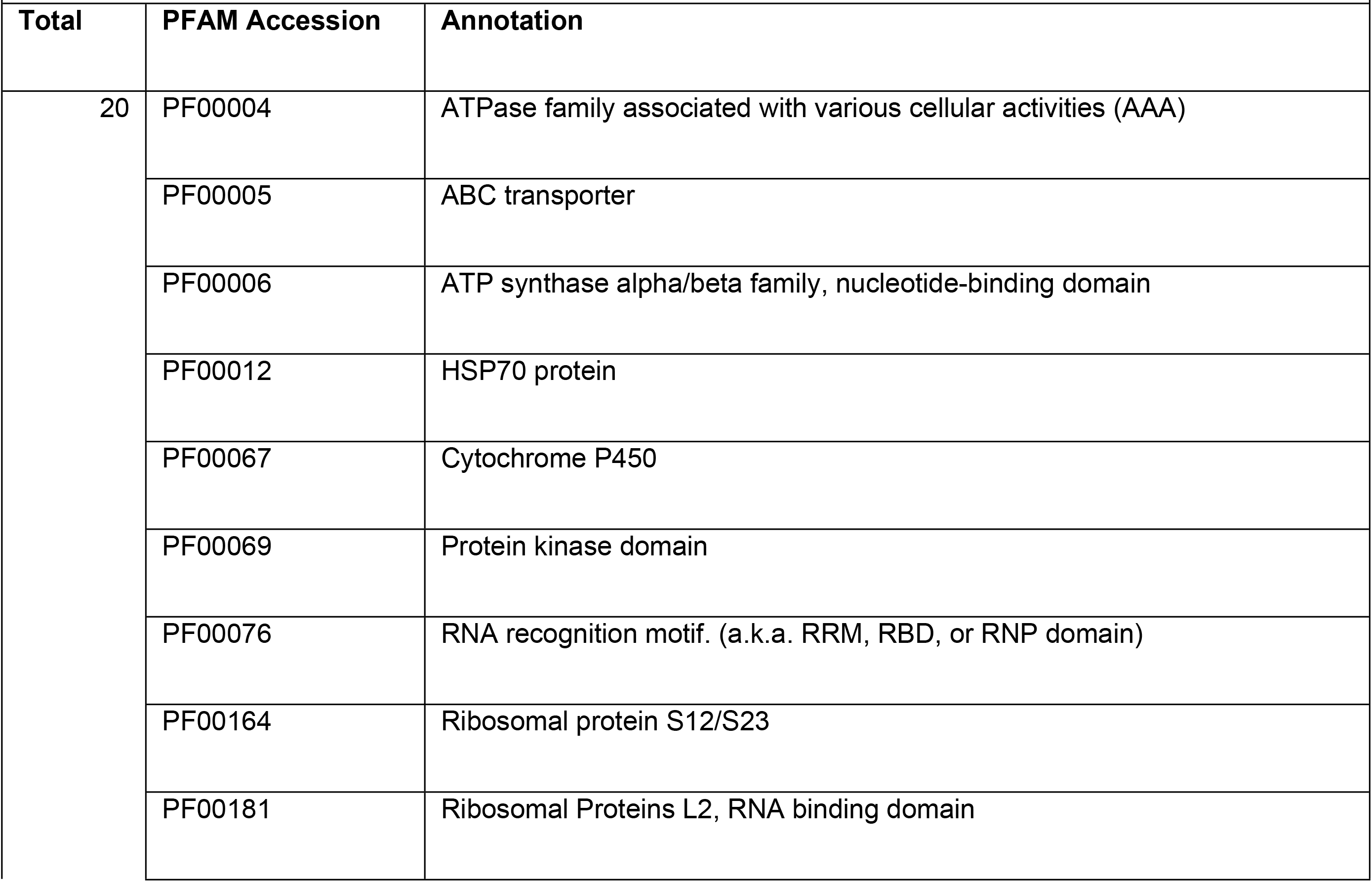
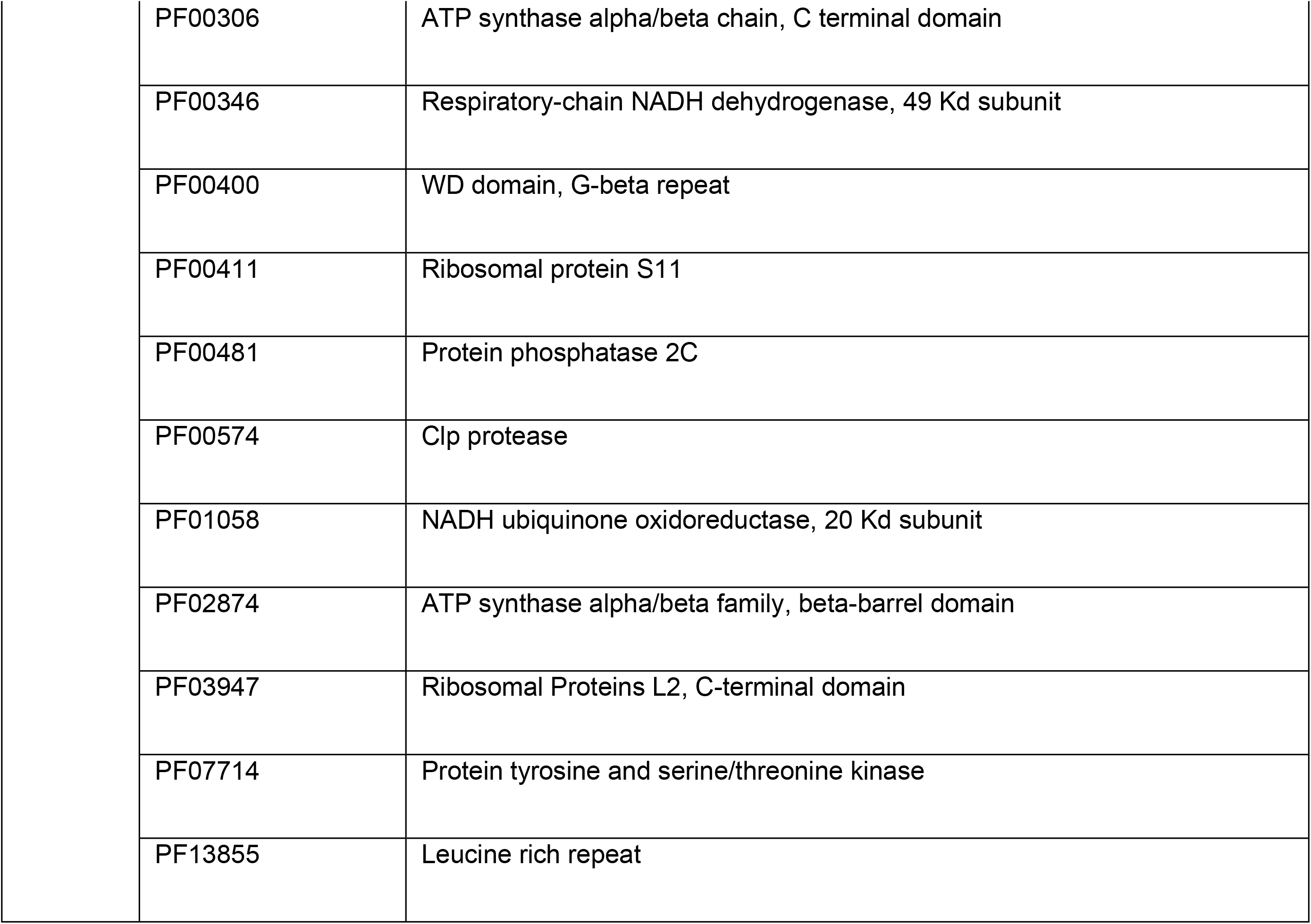
Gene elements shared by all GR eccDNA samples.

| Total | PFAM Accession | Annotation |
| --- | --- | --- |
| 20 | PF00004 | ATPase family associated with various cellular activities (AAA) |
|  | PF00005 | ABC transporter |
|  | PF00006 | ATP synthase alpha/beta family, nucleotide-binding domain |
|  | PF00012 | HSP70 protein |
|  | PF00067 | Cytochrome P450 |
|  | PF00069 | Protein kinase domain |
|  | PF00076 | RNA recognition motif. (a.k.a. RRM, RBD, or RNP domain) |
|  | PF00164 | Ribosomal protein S12/S23 |
|  | PF00181 | Ribosomal Proteins L2, RNA binding domain |

|  |  |
| --- | --- |
| PF00306 | ATP synthase alpha/beta chain, C terminal domain |
| PF00346 | Respiratory-chain NADH dehydrogenase, 49 Kd subunit |
| PF00400 | WD domain, G-beta repeat |
| PF00411 | Ribosomal protein S11 |
| PF00481 | Protein phosphatase 2C |
| PF00574 | Clp protease |
| PF01058 | NADH ubiquinone oxidoreductase, 20 Kd subunit |
| PF02874 | ATP synthase alpha/beta family, beta-barrel domain |
| PF03947 | Ribosomal Proteins L2, C-terminal domain |
| PF07714 | Protein tyrosine and serine/threonine kinase |
| PF13855 | Leucine rich repeat |

### Gene ontology enrichment of *A. palmeri* eccDNA

Gene ontology enrichment analysis of predicted coding elements on eccDNA of GS eccDNA revealed a variety of enriched biological processes, cellular components, and molecular functions encoded on eccDNA [Fig 4]. Enriched biological processes include regulation of transcription, membrane and lipid transport, DNA binding, fatty acid biosynthesis, protein phosphorylation, oxidation-reduction, chromatin maintenance, and protein translation [Fig 4a and S5 Table]. Cellular component and molecular function categories of interest include membrane and ribosome components [Fig 4b], cytoplasm, protein kinase activity, and ATP binding [Fig 4c].

**Fig 4.**
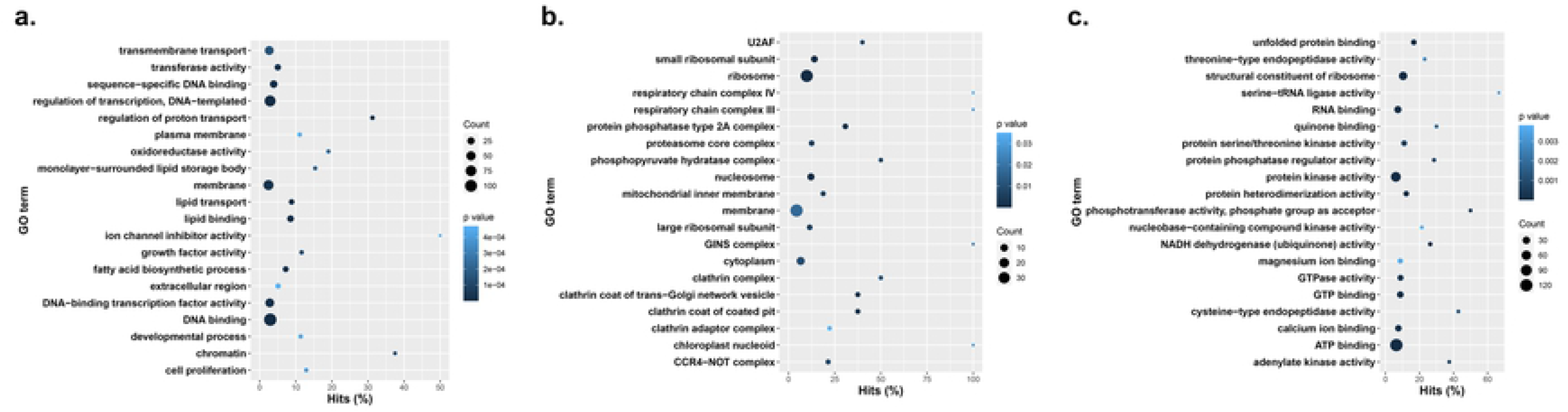
Gene ontology enrichment terms and their prevalence among GS *A. palmeri* eccDNA samples. **A.** Biological processes, **B.** cellular components, **C.** molecular functions.

Glyphosate resistant eccDNAs showed similar, but slightly different enriched biological processes such as transmembrane transport, translation, protein phosphorylation, and oxidation-reduction process [Fig 5a]. Ribosome, nucleus, membrane, and integral component of membrane were also enriched in the cellular component category [Fig 5b]. Representative molecular functions for GR eccDNA were mainly in the ribosome and membrane categories, but ATP binding, protein kinase activity, and catalytic activity were enriched [Fig 5c and S6 Table].

**Fig 5.**
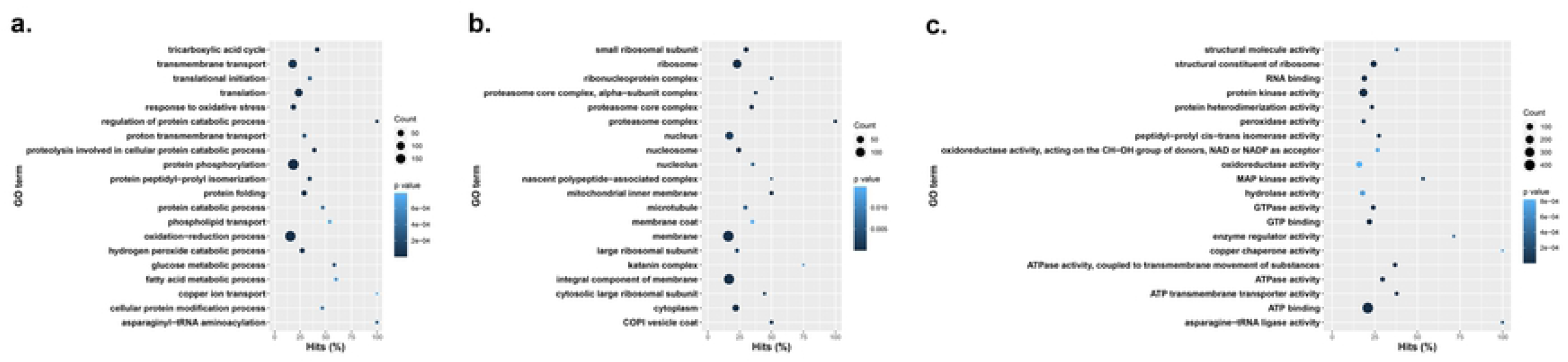
Gene ontology enrichment terms and their prevalence among GR *A. palmeri* eccDNA samples**. A**. Biological processes, **B**. cellular components, **C**. molecular functions.

### Repeat structure of *A. palmeri* eccDNA

Repeat characterization revealed a high proportion of repetitive sequences among both GS and GR eccDNAs [S7 Table]. The most common repeat classes were simple repeats, long terminal repeats (LTR) from the Copia superfamily, low complexity regions, and LTR from the Gypsy superfamily [S7 Table]. Interestingly, simple repeat content varied drastically among the GR and GS states. For example, Arizona and Mississippi GS and GR pairs were closely balanced in terms of content, but Mississippi has nearly 6 times as many with ∼17.5k compared to ∼4k simple repeats [S7 Table]. The Long Terminal Repeats/Copia class was second in abundance among eccDNAs, followed by low complexity repeats and then Gypsy elements. DNA elements such as Stowaway, LINES, Cassandra, hAT-Tip100, MULE-MuDR, and helitrons were also identified in both GS and GR biotypes [S7 Table].

### Similarity to the eccDNA replicon and replication origins on eccDNAs in *A. palmeri*

Alignment and comparative analysis for coding content and conserved sequence structure between GS and GR eccDNAs and the eccDNA replicon [26] identified a total of 162 GS eccDNA and 2,547 GR eccDNA with at matches at least 100 bp in length with a percent identify of at least 95% [Fig 6]. A total of 7 and 11 eccDNA replicon genes were predicted in GS and GR eccDNA, respectively [S8 Table]. Predicted eccDNA replicon genes in GS eccDNA include PEARLI4, Heat shock (HSP70), no apical meristem (NAM), replication factor-A, retrotransposon, zinc finger, and suppressor of gene silencing [S8 Table]. GR predicted replicon genes include: *EPSPS*, PEARLI4, Domain of unknown function (DUF), ethylene response factor, HSP70, NAM, replication factor A, and retrotransposon [S8 Table]. Interestingly, several GR eccDNA contained multiple copies of the EPSPS gene from Arizona, Delaware, Kansas, and Maryland, while the EPSPS gene was not present on any eccDNA in GS [Fig 7A].

**Fig 6.**
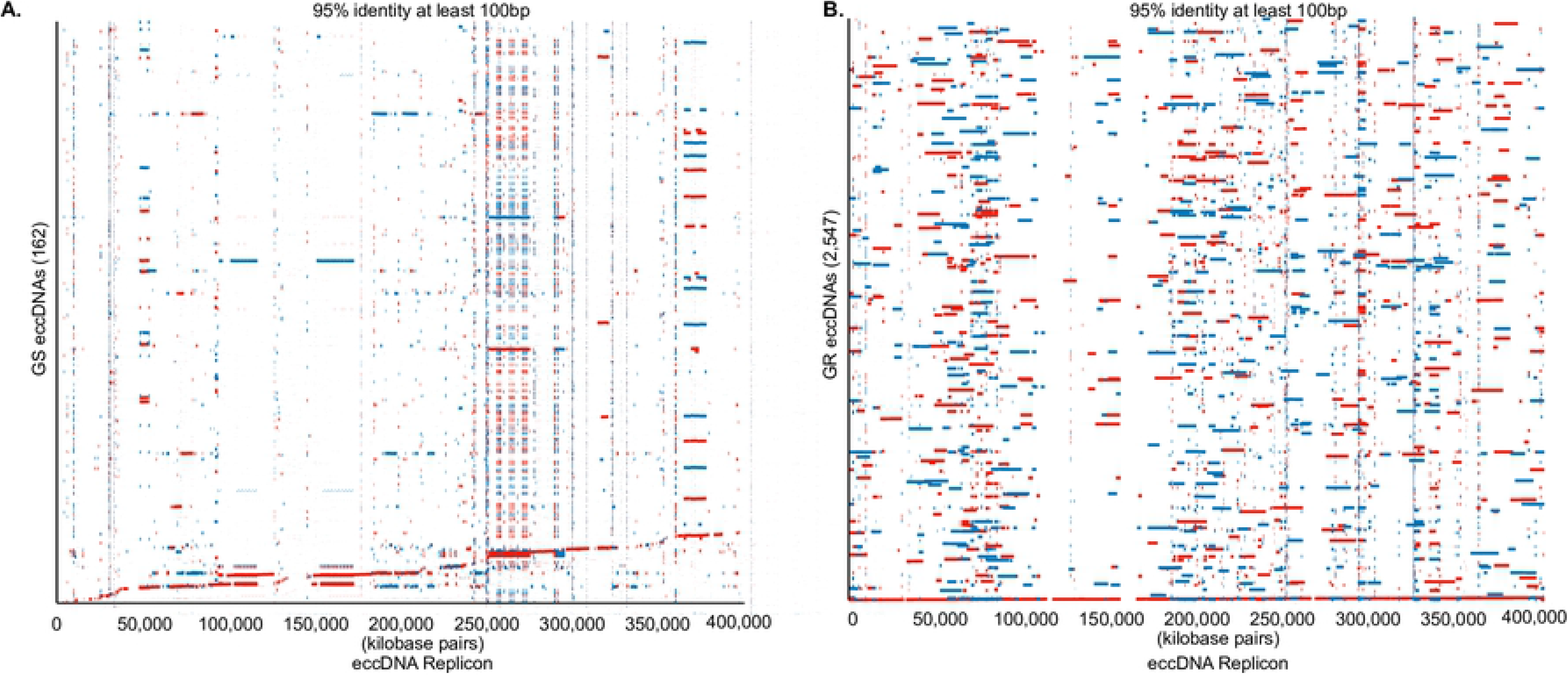
Alignment of eccDNA to the replicon in GS and GR biotypes. A. Alignment of 162 GS eccDNA to the eccDNA replicon. B. Alignment of 2,547 GR eccDNA to the eccDNA replicon. Red colors indicate indirect orientation and blue are direct. Alignments are filtered for matches of at least

**Fig 7.**
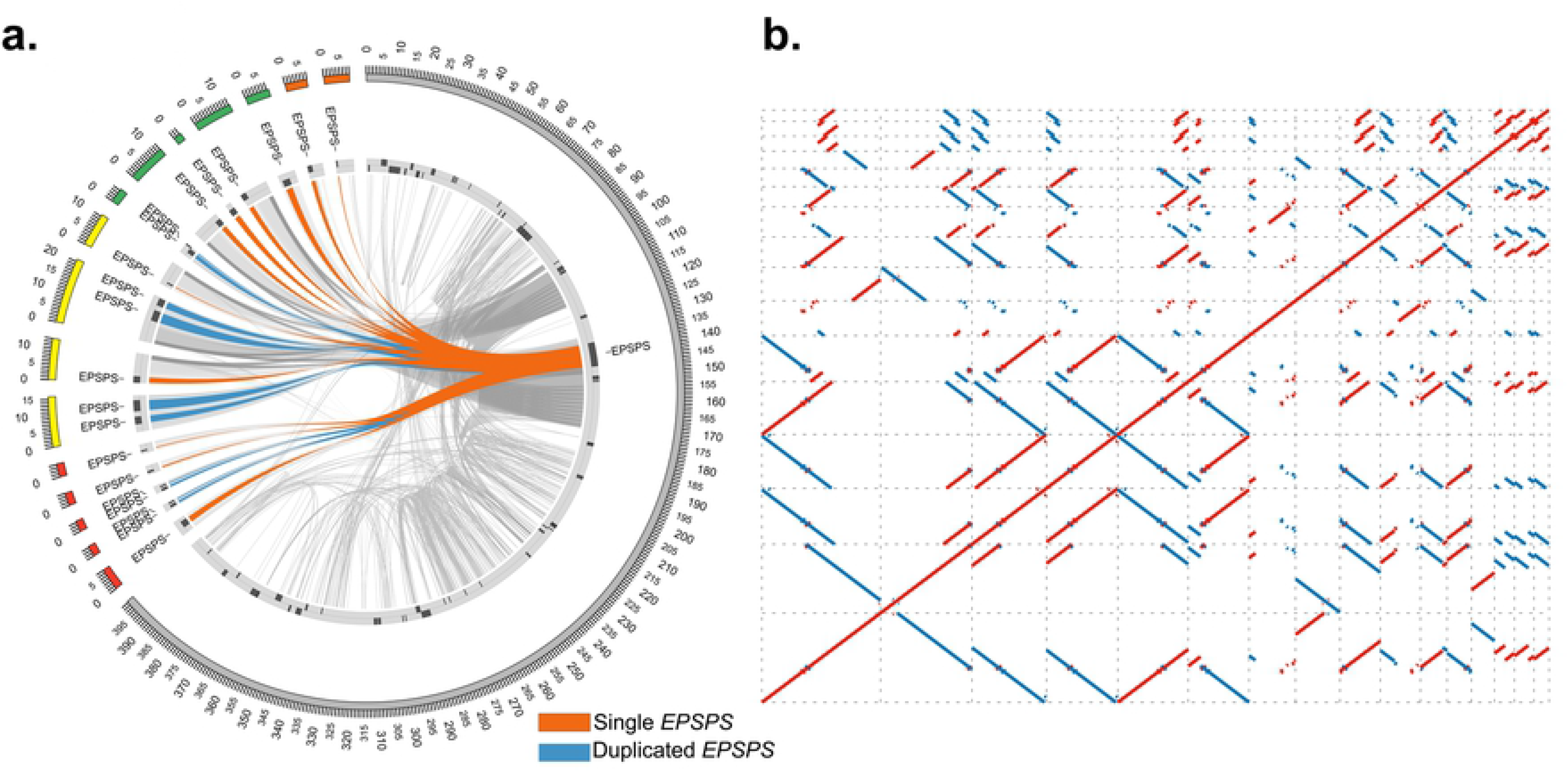
*EPSPS* gene copies in GR eccDNA. A. Sequence similarity of GR eccDNA aligned to the eccDNA replicon. Blue and orange links indicate single or duplicated EPSPS genes. Grey links show broader sequence similarities. B. Self-alignment of the GR eccDNA containing multiple EPSPS copies. Blue dots indicate inverted repeat sequences and red dots indicate repetitive sequence in the forward direction.

In GR eccDNA we identified 5 eccDNA with 2 copies of the *EPSPS* gene and 11 eccDNA with a single *EPSPS* copy [Fig 7A]. A self-alignment of the GR *EPSPS* eccDNA shows many conserved direct and indirect repeats [Fig 7B] with very high sequence identity (>95% with at least 100bp). Palindromic repeats that flank the *EPSPS* gene, previously described as possible genome tethering sites [26], were also evident among various eccDNA (Grey links in A and on the top right corner of B) indicating the potential for recombination among these smaller eccDNA, relative to the replicon.

Previous work has implicated a 17bp extended autonomous consensus sequence (EACS) with a motif of WWWWTTTAYRTTTWGTT that contains a core 11bp autonomous consensus sequence (ACS) reported in yeast [43] as a sequence where replication machinery initiates autonomous replication in plants [33]; which was functionally verified in the eccDNA replicon [35] [Fig 8]. Analysis of the GS and GR eccDNA for autonomous consensus (ACS) sequences (ACS) [43] identified a total of 430 unique eccDNA with 16 of the 17 bp present in the EACS with the common missing base being the first ‘W’ (A or T), several of which had multiple EACS sequences [Fig 8 and S9 Table]. A total of 36,237 core ACS sites (11bp) were predicted within 18,679 unique eccDNA out of the total 37,336 predicted eccDNAs implicating this sequence as a possible common origin of replication sequence among smaller eccDNA in *Amaranthus palmeri*. Of the eccDNA that contained ARS sequences, 2,785 were predicted to contain coding sequences, whereas 16,048 eccDNA did not contain an ARS sequence.

**Fig 8.**
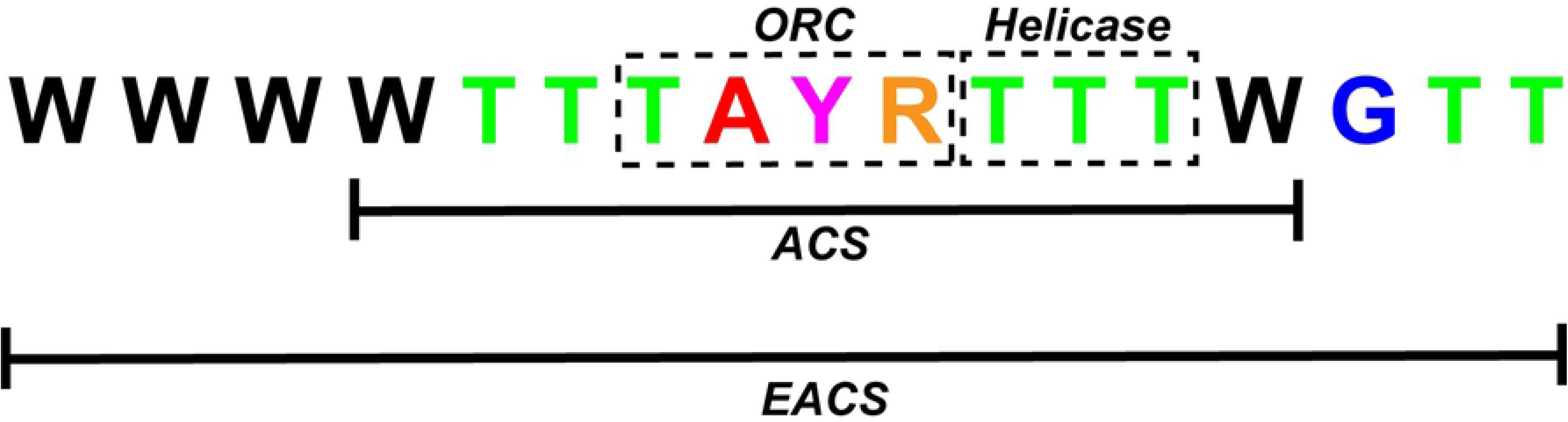
Extended autonomous consensus sequence sequence (EACS) presented in [34]. The core autonomous consensus sequence is highlighted with the TAYR motif highlighted as the origin of replication complex binding site (ORC) and the TTT motif highlighted as a helicase binding site. ‘W’ denotes A or T, ‘Y’ denotes C or T, ‘R’ denotes G or A.

### Genomic origins of eccDNAs in *A. palmeri*

To determine the genomic origins of eccDNA and the possibility of genomic regions with a disposition for eccDNA formation, GS and GR eccDNA were mapped to the chromosome scaffolded *Amaranthus palmeri* assembly [44] and counted using non-overlapping genomic windows of 500kb [Fig 9]. We identified several regions of the genome with a very high disposition for focal amplifications that are conserved between GS and GR. These regions include the distal end of chromosome 2 and near the center of chromosome 3, and several other regions distributed throughout the genome [Fig 9a]. The 500kb window localized at the distal end of chromosome 2 contained 285 eccDNA from GS and 469 from GR [Fig 9a and S10 Table]. The center of chromosome 3 contained 225 GS and 449 GR eccDNA. The genomic region of eccDNA origin among GR samples with the most eccDNA was on chromosome 4 with 487 eccDNA and only 51 from GS, suggesting a possible signal of glyphosate stress. Extraction and self-alignment of the 6 genomic windows from the Palmer amaranth chromosome scale assembly from [44] revealed intricate arrays of repetitive sequence [Fig 9b]. Short, inverted repeats were the most common among all 6 regions [Fig 9b]. Clustered palindromes of various sizes were discovered in segments 2, 3, 4, and 5, as indicated by box-like structures. Regions 2 and 3 (highlighted in Fig 9b) contained more complex repetitive structure with larger direct repeats (region 2) and indirect repeats (region 3) [Fig 9b].

**Fig 9.**
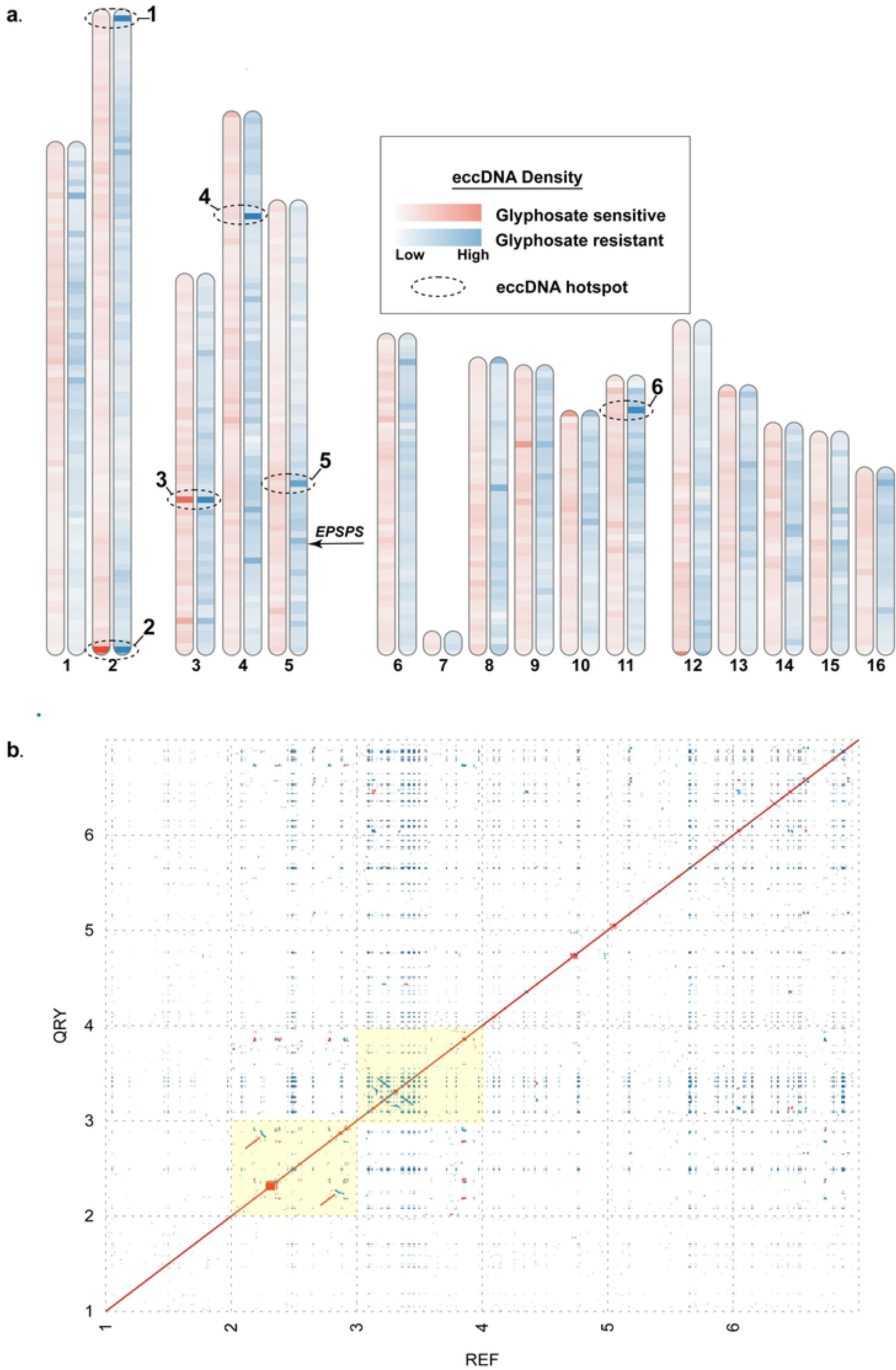
**A.** Alignment and quantification of unique eccDNA of GS (red) and GR (blue) to the chromosome-scale Palmer amaranthus reference assembly in 500kb non-overlapping windows. Darker colors represent a larger abundance of mapped eccDNA. Regions with the dotted ellipses indicate a high abundance of mapped eccDNA. **B.** Self-alignment of the 6 highlighted regions from A. Red dots indicated direct repeats, while blue are indirect. Regions highlighted in yellow are derived from the 2 genomic regions (2 & 3 - 500kb each) with the highest abundance of mapped eccDNA.

## Discussion

Gene copy number variation is a predominant mechanism by which organisms respond to selective pressures in nature. Focal amplifications of transcriptionally active chromatin as eccDNAs have been found in both abundance and diversity across higher and lower order eukaryotic species underpinning their importance as a vehicle for gene copy amplification. Advancements of single molecule sequencing and approaches to purify and directly sequence circular DNA have led to evidence that eccDNA may have a fundamental role in the cell and function and also function as a source of genetic heterogeneity in response to environmental pressures [1-4, 22, 26, 31, 42]. Previous work in *Palmer amaranth* demonstrated that several genes in addition to *EPSPS* were co-amplified on a large eccDNA (∼400kb) with sophisticated repetitive content and origins from distal segmental genomic regions [26]. This large eccDNA served as the vehicle for *EPSPS* gene copy amplification, but whether construction of this large eccDNA was the result of intramolecular recombination or recombination among a population of smaller eccDNA is unclear.

Using single molecule sequencing and the CIDER-Seq analytical pipeline [42], we identified diverse and abundant eccDNA species in both GS and GR biotypes collected from distal geographic regions that were previously reported [41]. The sizes of these eccDNA ranged from a few hundred base pairs to nearly 30kb in both biotypes and between 6 and 20% were predicted to contain genes which indicates that eccDNAs are present in *Amaranthus palmeri* without glyphosate exposure.

Gene enrichment analysis of both GS and GR eccDNA provided insight on biological processes and molecular functions enriched for activities related to a generalized stress response or important for rapid adaptation such as transcription regulation, development, chromatin, protein phosphorylation, oxidation-reduction, ribosomal and membrane components, protein kinase activity, and ATP binding. This indicates that eccDNAs may have a role in preserving important protein synthesis genes. Notably, transfer RNAs (tRNA) were predicted to reside on eccDNA in bothy GS and GR samples, but only on eccDNA that do not contain coding sequences. This was also shown by Wang et al., 2021 in Arabidopsis [19] and suggests that protein synthesis is a key attribute or component of the early response to stress and or the adaptive response. This finding also suggests that regulation of protein synthesis is perhaps as driven by eccDNA is an independent component of selection and directed gene focal amplifications as eccDNA. Furthermore, plants likely require additional copies of these protein synthesis genes for stress responses to produce significant immunity or defense products, as is the case for GR *A. palmeri* [1]. For example, transmembrane transport has been shown to plays an important role in adaptation of *Arabidopsis* to metalliferous soils [45], resource allocation and sensing under plant abiotic stress [46–48] and were enriched on GS *Palmer amaranth* eccDNA. Fatty acid biosynthesis is another category of enriched genes on GS eccDNA which has been implicated in signaling and plant defense to pathogens [49, 50].

At the gene level, there were a core set of 9 functional protein coding domains in common among the GS samples. Ribosomal proteins (circular rDNA), which are commonly reported as functional genes among eccDNA, were found among all 9 GS samples suggesting a common role for rDNAs as circular structures in plants [6, 31, 51, 52]. Interestingly, Cytochrome p450 and Clp protease domains were also present in each of the GS samples. Cytochrome p450s are a superfamily of genes that perform a suite of functions in plant development and protection from various stresses via multiple biosynthetic and detoxification pathways. Cytochrome p450 activity plays a central role detoxification of xenobiotics in various weed species [53–56], biosynthesis of hormones, fatty acids, sterols, cell wall components, biopolymers, and various defense compounds [57]. Clp proteases are proteolytic enzymes whose increased expression also play a protective role for the plant in both abiotic and biotic stress [58–60]. Clp proteases help maintain protein homeostasis in chloroplasts and remove nonfunctional proteins, which is essential during stress episodes when proteins tend to be more vulnerable to damage [20–22]. These core genes encoded on GS eccDNA may contribute Palmer amaranth’s innate ability to rapidly adapt.

GS biotypes shared the same 9 core functional domains as GS biotypes including Cytochrome p450 and Clp protease, in addition to 11 other domains indicating that eccDNAs are dynamic and their presence and coding structure may be the result of selective pressures. Notably, the additional functional domains in GR biotypes include additional ribosomal motifs, ABC transporters, HSP70 proteins, and leucine rich repeat (LRR) domains. ABC transporters are important for detoxification, environmental stresses and pathogen resistance [23] and may play a complementary role in glyphosate detoxification in addition to *EPSP* synthase over accumulation. The most abundant functional domain and conserved among all the samples is the HSP70 domain; which functions in protein maintenance and a wide variety of stress response mechanisms such as response to high temperatures [61], and was also a predicted gene on the eccDNA replicon [26]. Hsp70 have been reported to function by holding together protein substrates to help in movement, regulation, and prevent aggregation under physical and or chemical pressure in plants [61, 62] and have served as functional target in improving abiotic stress resilience in *Arabidopsis* [63] and other species. It is notable that the HSP70 is present in both GS and GR biotypes but is a core gene shared among all GR biotypes. The presence of Hsp70 on eccDNA suggests a possible role in glyphosate resistance, or perhaps, a genomic mechanism for rapid mitigation of heat and other abiotic stresses. Leucine rich repeat (LRR) domains are associated with protein-protein interactions, often as part of plant innate immune receptors [64]. Various transcription factors such as WRKY, bZIP, helicases, GATA (zinc finger), E2F, helix-loop-helix, TCP, and others were also predicted on *A. palmeri* eccDNA. Since transcription factor access to heterochromatin is limited by its compact structure, eccDNAs may provide a faster and more effective avenue for protein synthesis. Cancer cells with oncogenes encoded on eccDNAs appear to produce significantly more transcript copies compared to the same oncogenes encoded on linear DNA structures [14].

A primary question underlying the origins and structural dynamics of the large eccDNA replicon (∼400kb) [25, 26] is the mechanism by which it is assembled. The most likely scenarios are long-range genomic interactions and a compounded building event over short evolutionary time; or intramolecular recombination between smaller eccDNA with newly selected genomic focal amplifications resulting from glyphosate stress to form the larger structure, again over short evolutionary time scales. Here we show a moderate degree of eccDNA replicon coverage with GS eccDNA [Fig 6a], however there are large, disconnected gaps in coverage. It is notable that the *EPSPS* gene was not found on any GS biotype eccDNA in this study, while several other replicon genes were. One of the primary drawbacks to the CIDER-Seq methodology is the limitation of eccDNA size to the read length of the Pacific Biosciences Sequel II instrument [42] which means eccDNAs larger than an average read length will not be sequenced intact, such as the eccDNA replicon [26]. This limitation prevented the complete assembly of the *EPSPS* replicon, however the *EPSPS* gene and most other predicted eccDNA replicon genes were found in GR biotype eccDNA and coverage of the replicon was practically complete, with only a few small gaps. Furthermore, the *EPSPS* gene was found on smaller eccDNA in GR biotypes in multiple copies, which corroborates the work of Koo et al., that observed the extra-chromosomal *EPSPS* gene vehicle as multi-meric forms. [25]. Together, these results suggest that eccDNA are present as a basal source of genetic heterogeneity or rapid response mechanism, are selectively amplified, and the large eccDNA structure reported to confer glyphosate resistance is likely built by recombination among smaller eccDNA over rapid evolutionary timescales.

Another important observation and similarity with the eccDNA replicon are the high abundance and seemingly random distribution of the core 11bp autonomous consensus sequence and a longer more conserved 16bp extended autonomous consensus sequence [43] among approximately half of the GS and GR eccDNA. The greater abundance seems to be on eccDNA without coding sequences. These sequences were previously verified to function in autonomous replication and may be regulated mechanism, perhaps epigenetic or other, to maintain gene copy numbers in *A. palmeri*. In the eccDNA replicon, there is a single copy of the 17bp consensus sequence and 46 copies of the 11bp sequence, seemingly randomly distributed among the replicon [26, 35]. This observation further supports the possibility that the eccDNA replicon is the result of recombination among smaller eccDNA. It is also possible that there are alternate mechanisms or origins of replication on eccDNA in *A. palmeri* that are used to maintain and amplify copy number. Previous work showed that the coding components of the eccDNA replicon seem to be derived from distal regions of the genome [26], and evidence presented here show that eccDNA in both GS and GR seem to originate from all over the genome, Fig 9a. Here, we also demonstrate that there are segments of the genome, or perhaps a genomic context, with a disposition for focal amplifications. These genomic ‘hotspots’ are comprised of various repeat structures that may have facilitate eccDNA formation. There are also regions of the genome that seem to be activated as ‘hotspots’ in response to glyphosate stress that suggests eccDNA formation may also be a directed event, rather than random. It is still unclear if genes need to be in the ‘right’ genomic context for a focal amplification to occur, or if other regulatory/initiation mechanisms exist. This work provides evidence that eccDNA are a basal component of the cell and likely function as a reservoir of genetic heterogeneity in *A. palmeri* as part of the rapid adaptation program.

## Materials and methods

### Plant material and genomic DNA extraction

Seeds were collected from individual GR plants that had survived glyphosate application as previously described [30, 41]. Plants were grown in 9 × 9 × 9 cm plastic pots that contained a commercial potting mix (Metro-Mix 360; Sun Gro Horticulture, Bellevue, WA, USA). Seeds were sown on the potting mix surface and lightly covered with 2 mm of potting mix. Pots were sub-irrigated and maintained in a greenhouse set at a temperature regime of 30/25 ∘C (day/night) and a 15-h photoperiod under natural sunlight conditions supplemented with high-pressure sodium lights providing 400 μmol m−2 s−1. Sampling for whole genome sequencing was performed using a leaf from the third node of two representative plants from each population. Total DNA was extracted using a modified CTAB-based protocol with chloroform, isopropanol, and RNase A buffer [65]. Briefly, leaf material from each sample (approximately 20-100 mg) was ground into a fine powder using a mortar and pestle with liquid nitrogen, extracted with CTAB buffer, chloroform extracted, and ethanol precipitated. Total genomic DNA was resuspended in 50 μl of TE (10 mM Tris, 0.1 m MEDTA, pH 8.0) buffer containing RNaseA. The tube was incubated at 37°C for 30 minutes and stored at −20°C.

### EccDNA enrichment and sequencing (CIDER-seq)

Circular DNA enrichment sequencing (CIDER-Seq) was used to enrich, sequence, and analyze eccDNAs from the leaf tissue DNA extraction samples according to the protocol by Mehta et al., [17]. Because we wanted to survey the landscape of eccDNA, we did not perform a size exclusion step prior to enrichment. Otherwise, the circular DNA amplification, debranching reaction, and DNA branch release and repair stages closely followed the methods of Mehta et al., [42]. Enriched eccDNA for each sample [10] was individually barcoded following the manufacturer’s recommended protocol (Pacific Biosciences), pooled in equimolar amounts, and sequenced on a Sequel II single molecule sequencer (Pacific Biosciences).

### EccDNA sequence processing and analysis

Raw sequence reads were demultiplexed and circular consensus sequences analyzed with the SMRT link software (Pacific Biosciences). Parameters for CCS analysis were stringent and include: 1) predicted quality = 0.999; and 2) minimum read length = 1,000 bp. Processed reads were stored as .fastq files. Processed fastq files were analyzed with the packaged CIDER-seq software using the suggested approach to identify circular DNA. Predicted eccDNA were matched to the *A. palmeri* reference genome by Montgomery et al., [44]. After processing of predicted eccDNA, shorter duplicate eccDNAs were collapsed into the longest reference eccDNA with the CDhit software [66] with an identity threshold of 90%. Reference eccDNA were annotated for genuine open reading frames using the MAKER annotation pipeline [67] and evidence for genes derived from the *A. palmeri* published annotation [44]. Alignments to the reference genome were performed with the Minimap2 software [68] and comparative genome alignments performed with Mummer 4.0 [69]. Transfer RNAs were determined with the tRNAscan-SE software with default settings [70]. The *A. palmeri* reference assembly from [44] was divided into non-overlapping windows of 500kb and mapped eccDNA counted with BedTools [71].

## Supporting information

**S1 Table.** Summary and functional annotation of glyphosate sensitive eccDNAs.

**S2 Table.** Summary and functional annotation of glyphosate resistant eccDNAs.

**S3 Table.** Venn diagram result summary for glyphosate sensitive eccDNA samples with annotations.

**S4 Table.** Venn diagram result summary for glyphosate resistant eccDNA samples with annotations.

**S5 Table.** Gene ontology enrichment of all glyphosate sensitive eccDNA genes classified as biological **process (BP), cellular component (CC), and molecular function (MF).**

**S6 Table.** Gene ontology enrichment of all glyphosate resistant eccDNA genes classified as biological process (BP), cellular component (CC), and molecular function (MF).

**S7 Table.** Repeat characterization of eccDNA in glyphosate sensitive and resistant samples.

**S8 Table.** Summary and functional annotation of predicted eccDNAs in glyphosate sensitive and resistant biotypes from different states.

**S9 Table.** eccDNA with predicted EACs or ACS sequence.

**S10 Table.** Counts of eccDNA mapping to the *A. palmeri* genome.

## Notes

### Competing Interest Statement

The authors have declared no competing interest.

